# Proteoglycan 4 (PRG4) expression and function in dry eye associated inflammation

**DOI:** 10.1101/2020.10.01.318576

**Authors:** Nikhil G. Menon, Ruchi Goyal, Carolina Lema, Paige S. Woods, Gregory D. Jay, Linda H. Shapiro, Rachel L. Redfern, Mallika Ghosh, Tannin A. Schmidt

## Abstract

**Purpose:** Dry eye disease (DED) affects hundreds of millions worldwide. Proteoglycan 4 (PRG4) has been shown to improve signs and symptoms of DED in humans. The objectives of this study were to characterize endogenous PRG4 expression by telomerase-immortalized human corneal epithelial (hTCEpi) cells, examine how exogenous recombinant human PRG4 (rhPRG4) modulates cytokine and chemokine secretion in response to TNFα and IL-1β, explore rhPRG4 as a potential substrate and/or inhibitor of MMP-9, and to understand how experimental dry (EDE) in mice affects PRG4 expression.

**Methods:** PRG4 secretion was quantified by Western blotting and PRG4 expression by immunocytochemistry. Cytokine/chemokine release was measured by ELISA, and MMP-9 inhibition was quantified using an MMP-9 inhibitor kit. EDE was induced in mice, and PRG4 was visualized by immunohistochemistry in the cornea and Western blotting in lacrimal gland lysate.

**Results:** hTCEpi cells synthesize and secrete PRG4 *in vitro*, which is inhibited by TNFα and IL-1β. TNFα and IL-1β significantly increased secretion of cytokine IL-6 and chemokines IL-8, IP-10, RANTES, and ENA-78, and several of these chemokines were downregulated after cotreatment with rhPRG4. Fluorescently-labelled rhPRG4 was internalized by hTCEpi cells. rhPRG4 was not digested by MMP-9 and inhibited *in vitro* activity of exogenous MMP-9 both in solution and in the presence of human tears. Finally, EDE decreased corneal and lacrimal gland expression of PRG4.

**Conclusions:** These results demonstrate rhPRG4’s anti-inflammatory properties in the corneal epithelium and its contribution to ocular surface homeostasis, furthering our understanding of PRG4’s immunomodulatory properties in the context of DED inflammation.

## 1. Introduction

Dry eye disease (DED) affects hundreds of millions of people worldwide, and effective treatment options are lacking. It is associated with significant pain, limitations in daily activities, diminished vitality, poor health in general, and often depression.[1–3]. DED is very prevalent, affecting 10-20% of the population in those aged between 20 and 40 years and more than 30% of those above 70 years [2]. While treatment options do exist for DED, treatment options are often misaligned with the fundamental mechanisms of the disease and do not fully address patient symptoms [4]. Indeed, a recent study demonstrated more than 60% of DED patients using either of the two FDA-approved drugs had discontinued treatment within 12 months, thus indicating effective treatment options are lacking [5].

Recently, topical administration of a full length recombinant human proteoglycan 4 (rhPRG4) was shown to be clinically effective in improving signs and symptoms of patients with DED [6]. This small, single site, clinical trial (NCT02507934) on 39 subjects with moderate DED assessed the safety and efficacy of rhPRG4 at 150 μg/ml as compared to a 0.18% sodium hyaluronate (HA) eye drop in subjects with moderate DED. The rhPRG4 solution demonstrated statistically significant effects compared to HA in: a) symptomatic improvements in foreign body sensation, sticky feeling, blurred vision, and photophobia in at least one eye b) improvement in objective signs of DED in corneal fluorescein staining, tear film break-up time, eyelid and conjunctival erythema, and daily mean instillations; and c) no treatment-related adverse events. However, there remains a significant need for basic science research on rhPRG4’s biological properties and its potential therapeutic mechanisms of action in treating DED.

Tear film instability and tear hyper-osmolarity contribute to a positive feedforward loop in DED, leading to ocular surface damage and inflammation. Environmental stress, aging, bacterial infection, and contact lens wear can cause damage in the eye, specifically in the corneal epithelium and conjunctiva, which can lead to changes in tear film stability and osmolarity [7]. These changes can increase the production of pro-inflammatory cytokines, including interleukin (IL)-1β and tumor necrosis factor (TNF)-α, from corneal epithelial cells, which further degrades the tear film and contributing to a vicious inflammatory cycle [8,9]. Elevated levels of IL-1β, IL-6, and TNFα have also been reported in tears of patients with DED [10,11]. Changes in tear film stability and osmolarity also cause increased release of chemokines, including CCL5 (RANTES) [10,12], CXCL8 (IL-8) [11,13], and CXCL10 (IP-10), which attract immune cells to the corneal epithelium and further stimulate the proinflammatory cytokine production and matrix metalloproteinase (MMP) activity [7,8,12,13]. MMP-9 levels also increase in the tears of patients with DED, where they increase with DED severity, as well as in mice with experimental DED, and MMP-9 is used as a biomarker for DED [14–16]. Given the role inflammation plays in the pathogenesis of DED, molecules that are able to appropriately modulate inflammatory activity in the cornea could lead to potential, more effective, future treatment options.

PRG4, also known as lubricin, is a mucin like glycoprotein that is present on the ocular surface where it plays a critical protective role in maintaining ocular surface integrity. PRG4 is present at the epithelial surface of human cornea and conjunctiva, where it helps maintain the tear film [17]. Lack of PRG4 in mice results in increased ocular surface damage [18,19]. Mechanically, PRG4 functions as an ocular surface boundary lubricant, reducing friction at the interface between the eyelid, cornea, and contact lenses [20]. Full length rhPRG4 exhibits similar *in vitro* boundary lubricating properties to native PRG4, on both the ocular surface [19] and articular cartilage [21]. Biologically, recent studies in the context of synovial joint health and disease demonstrated that rhPRG4 has anti-inflammatory properties, including the ability to bind to and antagonize toll-like receptors, reducing inflammation by dampening nuclear factor kappa B (NFκB) activation and inflammatory cytokine expression [22], and the ability to inhibit fibroblast-like synoviocyte proliferation [23,24]. However, despite rhPRG4’s promising potential as a treatment for DED, rhPRG4’s anti-inflammatory properties and endogenous PRG4 expression in the context of ocular surface health and DED have yet to be examined.

The primary objectives of this study were therefore to characterize endogenous PRG4 expression by human corneal epithelial cells, and examine the ability of exogenous rhPRG4 to modulate cytokine and chemokine secretion in response to dry eye associated inflammatory stimuli, TNFα and IL-1β, in human corneal epithelial cells. Secondary objectives were to explore rhPRG4 as a potential substrate and/or inhibitor of MMP-9, and determine if EDE modulates corneal and lacrimal gland PRG4 expression.

## 2. Methods

### 2.2 Human Corneal Epithelial Cell Culture

#### 2.1.1 Expansion & Differentiation

hTCEpi cells (kindly provided by Dr. James Jester) [25] were expanded with Keratinocyte Serum-Free Medium (KSFM) supplemented with 5 ng/mL epidermal growth factor (EGF) and 25 μg/mL bovine pituitary extract (BPE) (Gibco, Grand Island, NY) and seeded in 12 well plates. Each well received 100,000 cells in 2 mL of KSFM media. Cells were allowed to expand, to 80-90% confluency, for 48 hours. Cells in each well were then differentiated using DMEM/F12 (Gibco, Grand Island, NY) + 10% FBS (R&D Systems, Minneapolis, MN) + 10 ng/mL EGF (Gibco), as described previously [26].

#### 2.2.2 Inflammatory Stimuli & rhPRG4 Treatment

After 4 days of differentiation, each well received 0.45mL of stock rhPRG4 (Lubris Biopharma, Framingham, MA) or sterile PBS containing 0.01% Tween-20 immediately followed by 1.55 mL of treatment media (DMEM/F12 + 12.9% FBS + 12.9 ng/mL EGF). The treatment media was spiked with IL-1β (Peprotech, Cranbury, NJ), TNFα (Peprotech) or left alone as a control. The final concentrations were 10 ng/mL for IL-1β, 100 ng/mL for TNFα, and 0 or 300 μg/mL for rhPRG4. After 48 hours of treatment, the conditioned media was collected, aliquoted, and stored at −20°C. Each treatment was tested in triplicate in each experiment, and the experiment was performed three times (N=3).

### 2.3 Western blot

Sodium dodecyl sulphate-polyacrylamide gel electrophoresis (SDS-PAGE) was performed with 3-8% Tris-Acetate gels (Invitrogen, Carlsbad, CA), as described previously [27]. Briefly, samples (15 μL of conditioned media per well) were electrophoresed, followed by electroblotting to a PVDF membranes and blocked in 5% non-fat dry milk (Biorad, Hercules, CA) in Tris-buffered saline + 0.05% Tween-20 (TBST) for 1h at room temperature. Membranes were then probed with anti-PRG4 Ab LPN (1:1000, Invitrogen, PA3-118) in 3% non-fat dry milk in TBST overnight at 4°C. After washing with TBST, membranes were incubated with HRP conjugated anti-rabbit secondary antibody (1:2000, MilliporeSigma, Burlington, MA) for 1 hour and imaged on a G:Box Chemi XX9 imager (Syngene, Frederick, MD) using SuperSignal West Femto (Thermo Fisher Scientific, Waltham, MA). Resulting intensity of bands were quantified by densitometry with the Genetools (Syngene).

### 2.4 ELISA analysis

Conditioned media samples were analyzed using commercially available ELISAs as per manufacturer guidelines (BioLegend, San Diego, CA), at 2X and 8X dilutions for IL-6, IP-10, RANTES, and ENA-78, and at 8X and 32X dilution of IL-8. Resulting data was collected on a SpectraMax i3x plate reader (Molecular Devices, San Jose, CA), and concentrations were calculated using OD values within the standard curve range.

### 2.5 Immunofluorescence Imaging

hTCEpi cells were seeded at a density of 30,000 cells per well on sterile glass coverslips in 4-well dishes. Cells were differentiated and treated using FITC-tagged rhPRG4 as described above (2.2.2). After 2 days, conditioned media was collected, and cells were fixed with 0.5 mL of 4% paraformaldehyde for 20 minutes at room temperature, permeabilized with 0.5 mL of 0.1% Triton X-100 in PBS for 5 minutes at room temperature, and blocked with 5% normal goat serum (Rockland, Pottstown, PA) and 5% BSA (MilliporeSigma) in PBS. Wells without FITC-rhPRG4 were probed with anti PRG4 Ab LPN (1:500, Invitrogen) in blocking solution overnight at 4°C. The next day, wells were appropriately treated with anti-rabbit 594 antibody (1:1200, MilliporeSigma) for 1 hour at room temperature and then DAPI (1:1000, Thermo Fisher Scientific) for 15 minutes at room temperature. The cover slips were then mounted with gold anti-fade mounting media (Thermo Fisher Scientific) and imaged on a Zeiss Axioskop2 microscope with an AxioCam digital camera (Zeiss, Oberkochen, Germany) using 63x oil objective. Slides with samples containing FITC-PRG4 were also imaged on a Zeiss LSM 880 confocal microscope with a 63x oil objective and a slice height of 0.36 μm.

### 2.6 MMP-9 Digestion & Inhibition

First, rhPRG4 degradation from exogenous MMP-9 was assessed using gel electrophoresis and SimplyBlue protein stain. Briefly, rhPRG4 at 0.77 mg/mL was incubated with or without 9 mU of MMP-9 enzyme for 60 minutes at 37°C. These samples were then electrophoresed on a 3-8% Tris-Acetate gel and stained with SimplyBlue, following manufacturer guidelines. Second, to assess the extent to which rhPRG4 inhibited *in vitro* MMP-9 activity, an MMP-9 inhibitor testing kit was used following manufacturer guidelines (Abcam, Cambridge, MA). This experiment was repeated using the same concentrations of rhPRG4 and MMP-9 in the presence of human tears. OD values were measured using an excitation wavelength of 328 nm and an emission wavelength of 420 nm on a SpectraMax i3x microplate reader.

### 2.7 Experimental Dry Eye Model

#### 2.7.1 Sample Collection and Preparation

Mice with experimental dry eye (EDE) were obtained as previously described [28,29]. Briefly, 8 to 12-week old C57BL/6 mice (equal number male and female) were housed in a controlled room in which humidity was maintained at approximately 20% and temperature was maintained at 21°C to promote ocular surface desiccation. Additionally, mice were given subcutaneous scopolamine hydrobromide injections three times daily for five consecutive days to reduce tear production. After treatment, whole eyes were collected, snap frozen in optimal cutting temperature compound to obtain frozen tissue sections, and stored at −80°C until PRG4 immunohistochemistry analysis. Lacrimal glands were also harvested and homogenized in 0.2% Triton X-100 (MilliporeSigma) containing a protease inhibitor cocktail (Roche) and prepared for PRG4 western blotting (2.2). Animal experiments were approved by the Institutional Animal Care and Use Committee at the University of Houston and adhered to the standards of the Association for Research in Vision and Ophthalmology Statement for the use of animals in ophthalmic and visual research.

#### 2.7.2 PRG4 Immunohistochemistry

Ten μm thick frozen eyeball sections were obtained on glass slides, fixed in cold acetone and then permeabilized with 0.1% Triton X-100/PBS. The sections were blocked with 15% goat serum (Abcam, Cambridge, United Kingdom) and then incubated overnight with anti-PRG4 Ab LPN primary antibody (Invitrogen) at 4°C. The sections were then incubated for 1h with Alexa Fluor 488 antibody (Abcam) and counterstained with DAPI. Images were captured by using the Delta Vision microscope (GE Healthcare, Chicago, IL). Fluorescence intensity was quantitated from captured images using ImageJ.

### 2.8 Statistical Analysis

Data are expressed as the mean ± SEM. The effect of inflammatory stimuli on endogenous PRG4 secretion by hTCEpi cells was assessed by one-way ANOVA followed by Dunnett’s post hoc testing. The effect of inflammatory stimuli on cytokine and chemokine secretion by hTCEpi cells was also assessed by one-way ANOVA followed by Dunnett’s post hoc testing. The effect of rhPRG4 treatment on cytokine and chemokine secretion within each inflammatory stimulus condition was assessed by one-tailed t-test. The effect of PRG4 on MMP-9 activity was assessed by one-way ANOVA followed by Tukey post-hoc testing. The effect of EDE on PRG4 localization in the cornea was assessed by one-tailed t-test. Finally, the effect of EDE on PRG4 expression in lacrimal gland lysate was assessed by one-tailed t-test.

## 3. Results

### 3.1 PRG4 secretion

First, to determine endogenous secretion of PRG4 as well as any changes with the addition of inflammatory stimuli, conditioned media from cells treated for 48h was collected and analysed using Western blotting. hTCEpi cells secreted PRG4 and this was reduced by treatment with inflammatory stimuli IL-1β and TNFα. Conditioned media from hTCEpi cells contained PRG4, in both monomeric and dimeric form, as assessed by western blotting (**Fig. 1A**). The addition of either IL-1β or TNFα reduced levels of PRG4 secretion to 0.84 ± 0.02 (p < 0.05) and 0.82 ± 0.06 (p < 0.05) fold of control, respectively (**Fig. 1B**).

**Figure 1.**
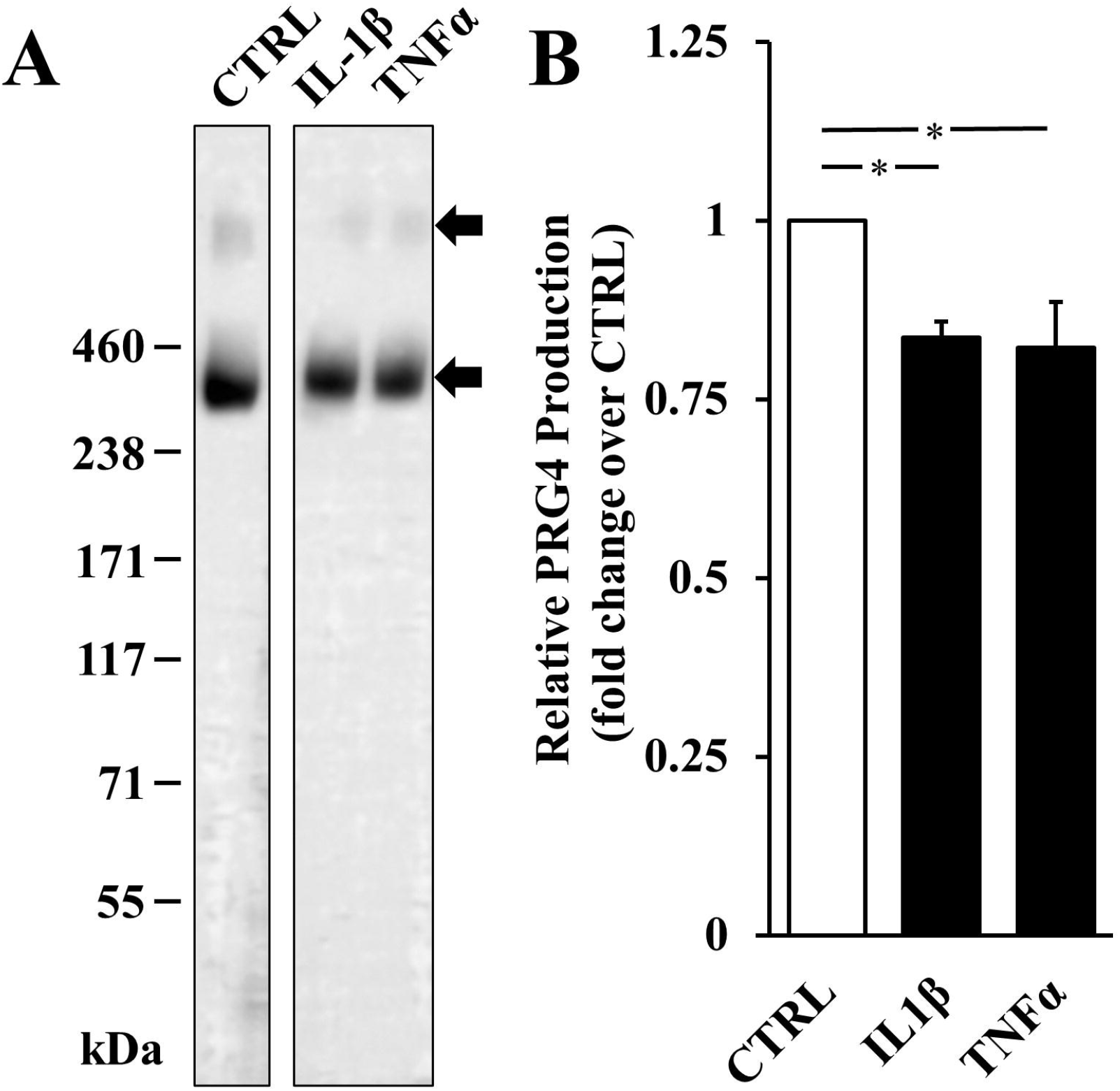
Effect of inflammatory stimuli on PRG4 secretion in hTCEpi cells. Western blot analysis of conditioned media samples from control, 100 ng/mL TNFα, or 10 ng/mL IL-1β conditions with anti-PRG4 Ab (A) densitometry of relative secretion compared to control (B). * = p < 0.05. Data are mean ± SEM (n=4)

### 3.2 Cytokine & Chemokine Secretion

To determine how cell secretion of cytokines and chemokines are influenced by inflammatory stimuli and rhPRG4, cells were treated with IL-1β or TNFα with or without the addition of rhPRG4 for 48h, and conditioned media was collected and analysed by ELISA. IL-1β and TNFα increased inflammatory cytokine and chemokine production by hTCEpi cells. TNFα significantly increased production by hTCEpi cells for all five cytokines and chemokines analysed: 15.02 ± 6.56-fold for IL-6 (from 1.32 ± 0.33 ng/mL, p < 0.05), 9.69 ± 3.70-fold for IL-8 (from 23.29 ± 10.40 ng/mL, p < 0.01), 3.72 ± 0.77-fold for IP-10 (from 329.40 ± 74.31 pg/mL, p < 0.01), 3.9 ± 1.18-fold for RANTES (from 28.19 ± 3.05 pg/mL, p < 0.001), and 5.30 ± 1.17-fold for ENA-78 (from 44.80 ± 18.14 pg/mL, p < 0.001) (**Fig. 2**). IL-1β only provided a significant increase of 10.52 ± 4.49-fold for ENA-78 (from 44.8 pg/mL, p < 0.05) (**Fig. 2**).

**Figure 2.**
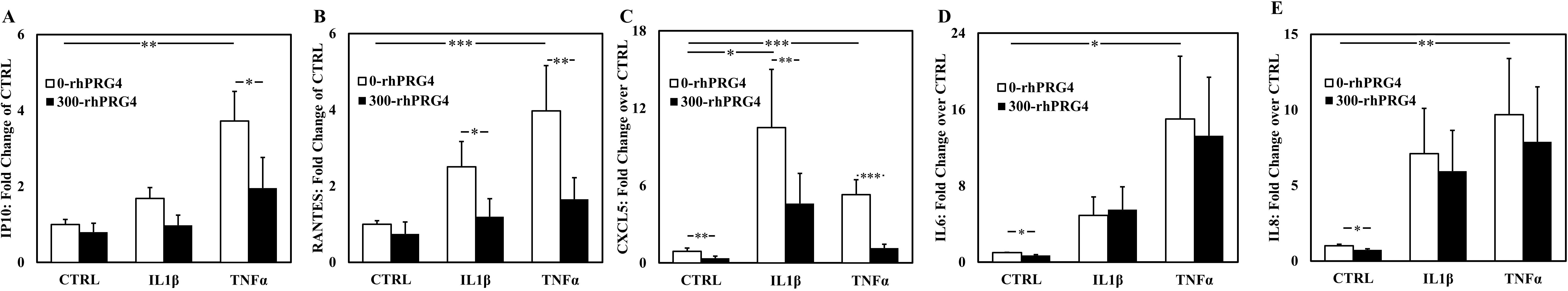
Effect of rhPRG4 on proinflammatory cytokine and chemokine secretion with inflammatory stimuli. Relative fold change in production of IL-6 (A), IL-8, (B), IP-10 (C), RANTES (D), ENA-78 (E) by differentiated hTCEpi cells treated with 100 ng/mL TNFα or 10 ng/mL IL-1β with or without 300 μg/mL of rhPRG4, measured by ELISA, * = p < .05, ** = p < .01, *** = p < .001. Data are mean ± SEM (n=10)

Treatment with rhPRG4 decreased production of several inflammatory cytokines/chemokines (**Fig 2**). In the absence of inflammatory stimuli, exogenous rhPRG4 significantly reduced levels of IL-6 (to 0.70 ± 0.08-fold, p < 0.01), IL-8 (to 0.73 ± 0.08-fold, p < 0.05), and ENA-78 (to 0.37 ± 0.17-fold, p < 0.05) in the conditioned media. For samples treated with IL-1β, exogenous rhPRG4 significantly reduced levels of RANTES (from 2.50 ± 0.67-fold to 1.20 ± 0.47-fold, p < 0.05) and ENA-78 (to 4.61 ± 2.35-fold, p < 0.05). For samples treated with TNFα, exogenous rhPRG4 significantly reduced levels of IP-10 (to 3.72 ± 0.77-fold, p < 0.05), RANTES (to 1.65 ± 0.57-fold, p < 0.01), and ENA-78 (to 1.15 ± 0.31-fold, p < 0.001).

### 3.3 PRG4 Visualization and Internalization

To visualize endogenous PRG4 expression and exogenous rhPRG4 internalization under normal and inflammatory conditions, hTCEpi cells were seeded onto glass coverslips and received IL-1β or TNFα with or without FITC-tagged rhPRG4 for 48h. Endogenous expression and rhPRG4 internalization was assessed by immunofluorescence and confocal microscopy, respectively. hTCEpi cells expressed PRG4 endogenously and were able to internalize exogenous rhPRG4 in response to inflammatory stimuli. Endogenous PRG4 was immunolocalized within and at the cell surface of hTCEpi cells both with and without the addition of inflammatory stimuli (**Fig. 3A-C**). Confocal imaging demonstrated slices within the cell contained exogenous FITC-tagged rhPRG4 as punctate staining (supplemental data), confirming that it can be internalized by hTCEpi cells in both control and IL-1β stimulation (**Fig 3D-F**). Interestingly, TNFα stimulation led to internalization of PRG4 expression with apparent loss of surface staining.

**Figure 3.**
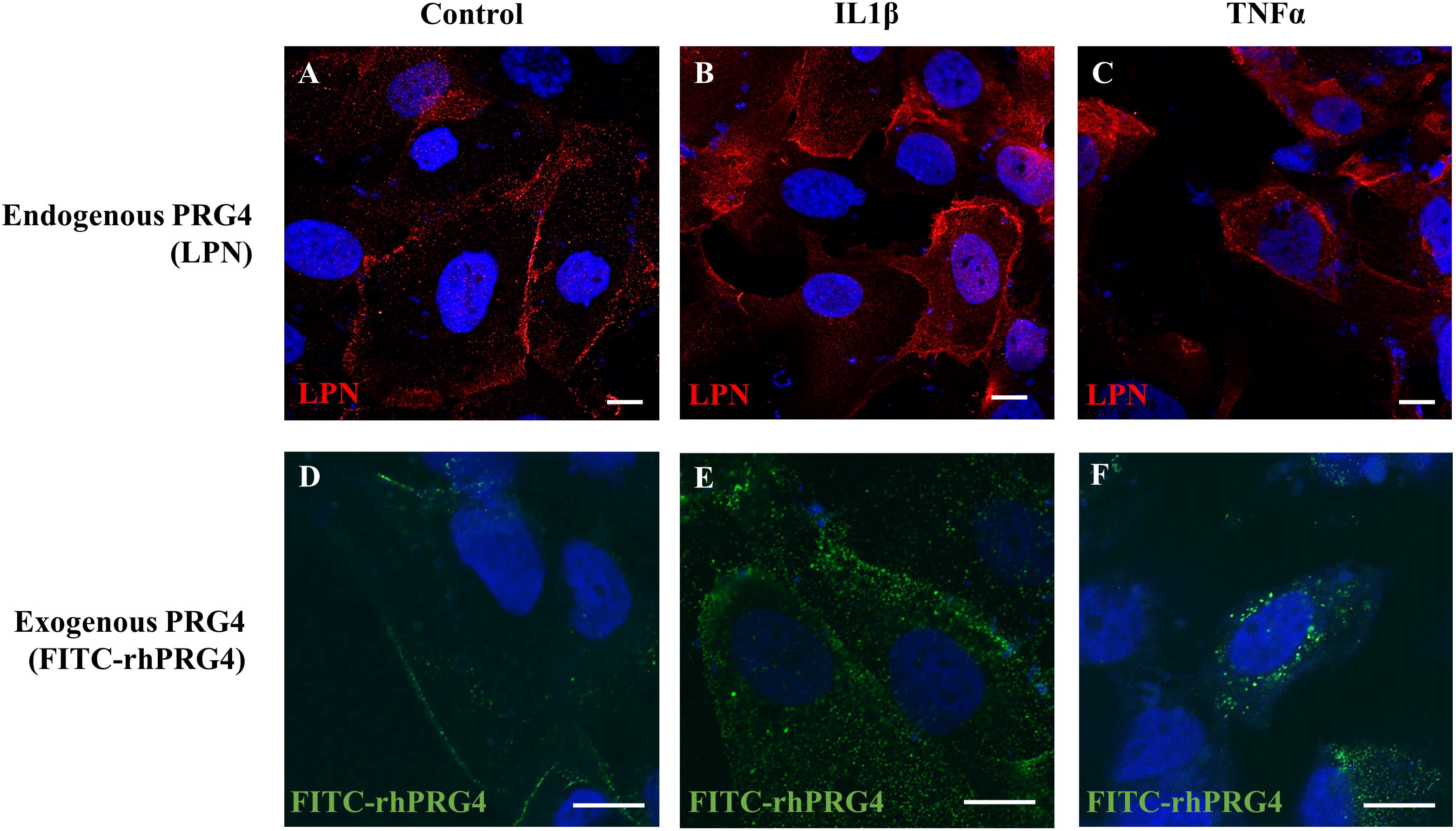
Effect of inflammatory stimuli on endogenous and exogenous PRG4 immunolocalization. Immunolocalization of PRG4 in hTCEpi cells with anti-PRG4 Ab LPN (A), and those treated with 10 ng/ml IL-1β (B) or 100 ng/ml TNFα (C). Confocal microscopy visualization of hTCEpi cells treated with FITC-tagged PRG4 (D), along with IL-1β (E), or TNFα (F). Nuclei are stained with DAPI. Scale bar is 10 μm, 63X magnification for A-C, 63X (x2) magnification for D-F.

### 3.4 Inhibition of MMP-9 activity by PRG4

rhPRG4 is not degraded by MMP-9, and *in vitro* activity of exogenous MMP-9 was inhibited by rhPRG4 both in solution and in the presence of human tear. No evidence of rhPRG4 *in vitro* degradation by exogenous MMP-9 was observed using gel electrophoresis and protein staining after co-incubation for 1h at 37°C (**Fig. 4A**). Analysis with an *in vitro* MMP-9 specific inhibitor assay demonstrated that rhPRG4 reduced *in vitro* activity of exogenous MMP-9 to 23% ± 11% for 150 μg/mL (p < 0.001), 16.5% ± 8.2% for 300 μg/mL (p < 0.001), and 5.3% ± 2.6% for 450 μg/mL (p < 0.001, **Fig. 4B**) compared to no rhPRG4 control. rhPRG4 also reduced *in vitro* activity of exogenous MMP-9 in the presence of human tears, (**Fig. 4C**) to 80.9% ± 6.6% for 150 μg/mL (p > 0.05), 41.1% ± 7.6% for 300 μg/mL (p < 0.001), and 15.6% ± 5.7% for 450 μg/mL (p < 0.001, **Fig. 4B**) compared to no rhPRG4 control. Additionally, there was a dose-dependent effect in reducing exogenous MMP-9 activity in the presence of tears (p < 0.001 – p < 0.05).

**Figure 4.**
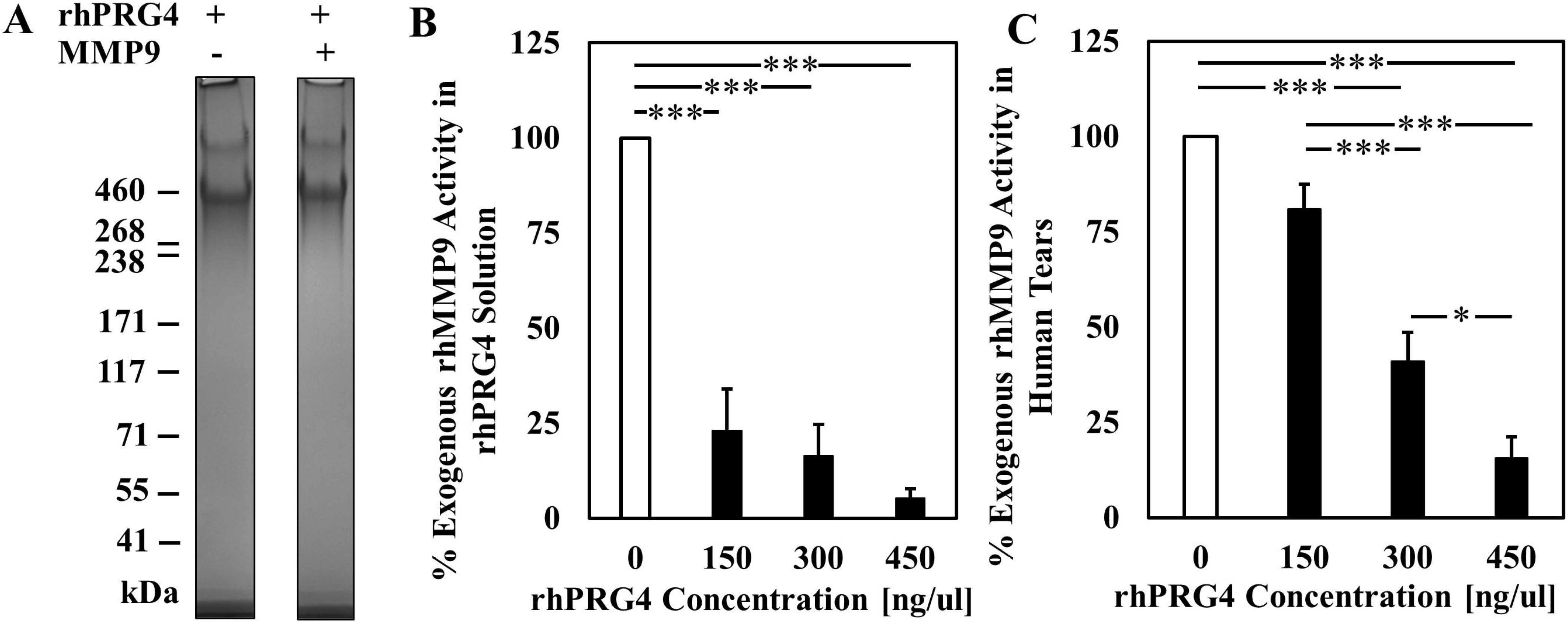
Effect of rhPRG4 on exogenous MMP9 activity *in vitro*. rhPRG4 incubated with or without MMP9 for 60 minutes at 37C (A). Exogenous MMP9 activity with 150 ug/mL, 300 ug/mL, and 450 ug/mL of rhPRG4 in solution (B) or in the presence of normal human tears (C). * = p < .05, *** = p < .001. Data are mean ± SEM (n=6)

### 3.5 Experimental dry eye model

To evaluate changes in PRG4 expression in an EDE model, mouse corneas from untreated and EDE mice were evaluated by IHC and lacrimal gland lysate samples were assayed by Western blotting. PRG4 expression was significantly reduced in both the cornea and lacrimal gland in EDE. PRG4 immunolocalization was clear in the cornea epithelium of untreated mice (**Fig. 5A**) and was reduced in the corneas of mice with EDE (**Fig. 5B**); there was a 2.96-fold reduction in the mean pixel intensity of PRG4 immunostaining between corneas of untreated mice and mice with EDE (**Fig. 5C**, p < 0.01). Lacrimal gland lysate from mice with EDE had reduced PRG4 production of 0.40 ± 0.24-fold compared to lysate from untreated mice (**Fig. 5D-E**, p < 0.05).

**Figure 5.**
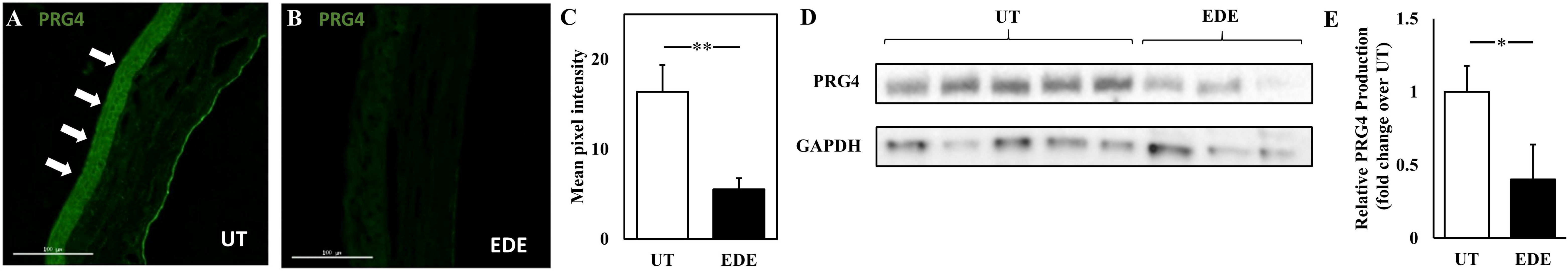
Effect of experimental dry eye (EDE) on PRG4 expression in the cornea and the lacrimal gland *in vivo*. PRG4 in mouse cornea immunolocalized with anti-PRG4 Ab LPN from untreated mice (A) and mice with EDE (B), quantified using ImageJ, * = p < .05, n = 4 (C). Lacrimal gland lysate analyzed by western blot using anti-PRG4 Ab LPN as primary antibody (D), Densitometry of relative fold change in PRG4 (E). ** = p < .01, Data are mean ± SEM (n = 3-5)

## 4. Discussion

The *in vivo* results of this study demonstrated for the first time that ocular surface PRG4 can be diminished in an experimental model of DED. In this model, which results in ocular surface damage and reduced tear production [28], PRG4 immunolocalization was reduced at the corneal epithelium, and PRG4 expression in lacrimal gland cell lysate was reduced. While unchallenged PRG4-deficient mice were previously shown to have increased ocular surface damage [18], the current data establish a relationship between DED and diminished PRG4 expression at the ocular surface. Given the critical role PRG4 seems to play in maintaining homeostasis, the reduction of PRG4 may be a core mechanism in the progression of DED, leading directly to increased evaporation, increased friction, and a pro-inflammatory environment. The epithelial glycocalyx, of which PRG4 is likely an integral part, is similarly involved in the retention of water at the ocular surface as the lipid layer. The data herein suggest that the loss of PRG4 expression in the corneal epithelia is at least a co-equal aspect of the positive feedback loop of disease pathogenesis in DED.

In addition, the *in vitro* results presented here extend previous studies examining PRG4 on the ocular surface as well as the regulation of PRG4 expression by other cell types. While PRG4 has previously been immunolocalized in human corneal epithelium [18], the results here demonstrate that PRG4 is expressed and secreted by hTCEpi cells and is downregulated by IL-1β and TNFα. This is also consistent with previous studies examining PRG4 secretion in synoviocytes [23], in chondrocytes with cartilage explants [30], and in synovial fluid after unilateral anterior cruciate ligament injury, where an increase in the concentration of inflammatory cytokines, including IL-1β and TNFα was associated with a reduction of PRG4 [31]. Given the parallels with other organ systems, these data suggest the addition of exogenous rhPRG4 may contribute to restoring homeostatic biological conditions [21,23,24,32,33].

A key finding of this study is that exogenous rhPRG4 can inhibit hTCEpi production of several chemokines stimulated by proinflammatory signals. Specifically, rhPRG4 treatment inhibited the RANTES, IP-10, and ENA-78 response to IL-1β or TNFα (**Fig. 2**). RANTES is known to play a key role in promoting lymphocyte migration to the corneal epithelium [34] and is upregulated in the tears of patients with DED [14]. IP-10 is a chemoattractant for Th1 lymphocytes and monocytes [35], and it is also upregulated in the tears of patients with DED [14]. ENA-78, which is involved in neutrophil activation and chemotaxis [36], is another chemokine found to be upregulated in the tears of patients with DED [37], and it has not been extensively studied in the corneal epithelium. While PRG4 purified from synovial fluid was previously shown to have anti-inflammatory effects in an induced-DED blinking eye model, where it reduced secretion of IL-1β, TNFα, and IL-8 [38], the present study demonstrates rhPRG4 has an *in vitro* anti-inflammatory effect on chemokines and long range signals to a variety of immune cell classes. Accordingly, the anti-inflammatory effect of PRG4 may stem from its ability to normalize the microenvironment through simultaneous friction reduction, downregulation of proinflammatory cytokines, and reduction of the recruitment of new immune cells. This pleiotropic nature of the molecule is unique, and strongly motivates its use for, and study in, DED, where multiple coincident etiologies are commonly observed.

Another key finding of this work is the visualization of exogenous FITC-labelled rhPRG4 being internalized by hTCEpi cells (**Fig. 3**). Both in basal conditions and with the addition of inflammatory stimuli, exogenous rhPRG4 was internalized and localized inside the cell, as visualized through confocal microscopy (**Fig. 3**), suggesting coupling of the molecule to intracellular signaling pathways, as seen in macrophages previously [39]. Given that rhPRG4 can modulate NFκB activity in other systems [24,38], future studies could also examine the effect of rhPRG4 on NFκB signaling, or other signaling pathways, as a potential mechanism by which rhPRG4 exerts its anti-inflammatory properties on corneal epithelial cells. Interestingly, and qualitatively, there appeared to be an increase in internalization with the addition of TNFα. Future work examining colocalization of rhPRG4 with endosomal markers will provide important clues to the fate of rhPRG4 post internalization. It is also worth noting rhPRG4 tagged with FITC also retained its ability to reduce levels of cytokine and chemokine secretion from hTCEpi cells (data not shown). This suggests that rhPRG4’s biological activity, at least in the context studied here, was not negatively affected by fluorescent tagging and therefore FITC-tagged rhPRG4 could be a useful reagent in the future to explore PRG4’s internalization into cells as well as its biological mechanism of action. rhPRG4 has been previously shown to be internalized in macrophages mediated, in part, through interactions with CD44 [39]. However, the precise processes by which PRG4 is internalized in hTCEpi cells (i.e. through receptor interactions, endocytosis, or a combination of the two) and the role it plays in rhPRG4’s anti-inflammatory mechanism of action remain under study.

Initial *in vitro* analysis here demonstrated that rhPRG4 is not a proteolytic substrate of MMP-9 (**Fig. 4**). While PRG4 is a substrate of other enzymes, including cathepsin S [40], cathepsin G [41], MMP-1 [42], and MMP-7 [42], it is not subject to degradation by MMP-9. The data also indicate that rhPRG4 inhibits *in vitro* activity of exogenous MMP-9 (**Fig 4**). Other studies have demonstrated that exogenous purified PRG4 can decrease MMP-9 levels in tears from a DED cell model [38] and MMP-9 gene expression stimulated by IL-1β in synoviocytes [23]. However, the inhibition of MMP-9 activity observed here is an important and potentially significant discovery. This inhibition effect was observed in solution and in the presence of human tears in a dose dependent manner. rhPRG4 may be inhibiting MMP-9 activity through the binding of the hemopexin-like (PEX) domain present on both rhPRG4 and MMP-9 [43,44]. The MMP-9 PEX domain binds to CD44, gelatin and a4ß1 integrins, and its inhibition may prevent homodimerization and reduce additional inflammatory signaling, including through MMP-9 damage-associated molecular pattern related pathways, such as Myd88 and TLR4, which are strongly associated with DED severity [28,45] Given the role MMP-9 plays in the pathogenesis of DED, these data may help elucidate some of the clinical efficacy of PRG4 in treating DED

## 5. Conclusion

In conclusion, these results underpin the understanding of PRG4 expression and anti-inflammatory properties of rhPRG4 within the context of corneal epithelium in DED. The demonstrated clinical utility of rhPRG4 in improving signs and symptoms in a small single site clinical trial, combined with ongoing research studying rhPRG4’s anti-inflammatory properties in other systems [32,39,46,47] reveals a therapeutic mechanism of action. Collectively these findings provide the foundation and motivation for continued basic research in further understanding PRG4’s properties on the ocular surface, both *in vitro* and *in vivo*, as well as expanding clinical evaluation of its ability to effectively provide relief to those who suffer from DED.

## Acknowledgments

We also gratefully acknowledge Dr. David Sullivan and Wendy Kam (Schepens Eye Research Institute, Boston MA) for aid with the culture of the hTCEpi as well as Dr. James Jester (University of California Irvine) for providing the cells. Finally, we thank Dr. Benjamin D. Sullivan (TearLab Corp) for discussions on and critique of the manuscript.

**Supplemental Figure**: hTCEpi cells treated with FITC-tagged PRG4 and no stimulus (A), IL-1β (B) or TNFα (C). Nuclei are stained with DAPI. Scale bar is 10 μm, 126X magnification

