## Supplementary figures and images for "Proteoglycan 4 (PRG4) expression and function in dry eye associated inflammation"

### Supplemental Figure 1

#### Slide 1
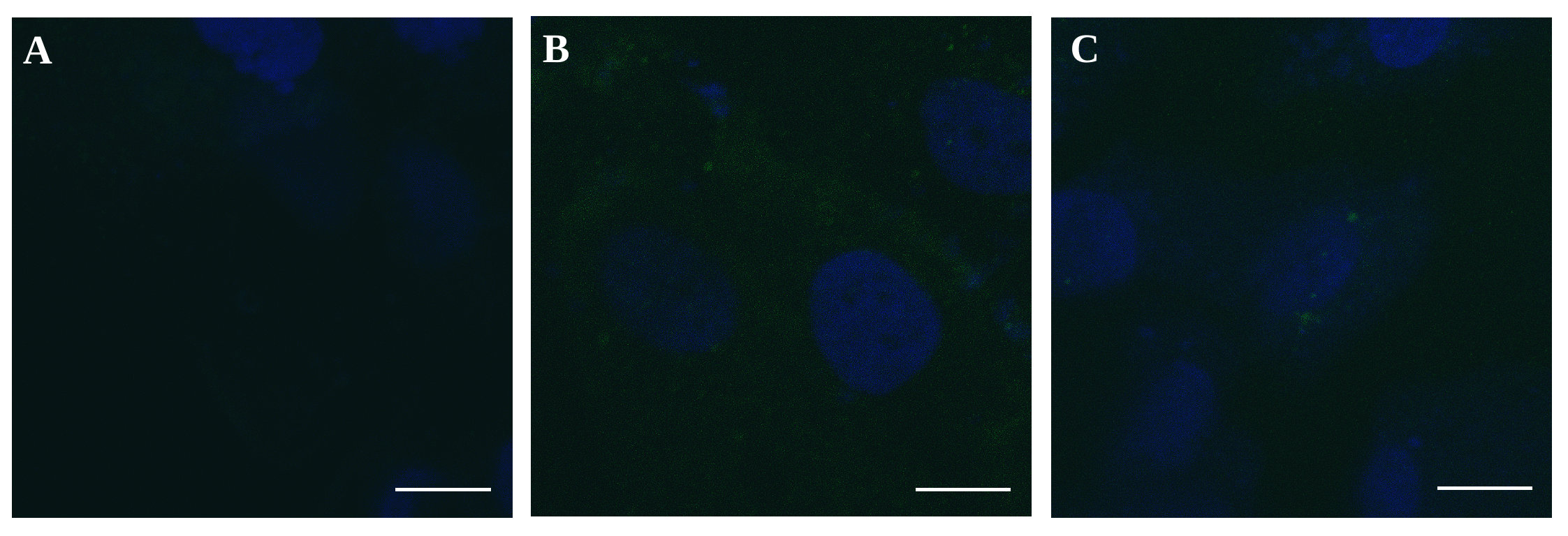

B
C
A
